# Porosome reconstitution therapy: A biologic rescue from cystic fibrosis

**DOI:** 10.1101/2024.09.11.612494

**Authors:** Cho Won Jin, Ha Vo, Yongxin Zhao, Douglas J. Taatjes, Bhanu P. Jena

## Abstract

Cystic fibrosis (CF) is a genetic disorder resulting from mutations in the CF Transmembrane Conductance Regulator (CFTR) gene that codes for a chloride transporting channel at the cell plasma membrane. In CF, highly viscous mucus is secreted in the airways preventing its clearance, leading to lung infections and respiratory failure. A major challenge in treating CF patients has been the presence of more than 2,000 different CFTR mutations or due to the absence of CFTR expression. CFTR is among the 34 major proteins composing the 100 nm porosome secretory machinery in the human airway epithelia, involved in mucin secretion. The airways is coated with a thin film of mucus, composed primarily of mucin MUC5AC and MUC5B. Sputum from patients with CF show a >70% decrease in MUC5B and MUC5AC secretion. Our studies using differentiated 3D cultures of human airway epithelial cell line, also demonstrate loss of both chloride and mucus secretion following exposure to CFTR inhibitors thiazolidinone 172 or the hydrazide GlyH101. Our studies show that human bronchial epithelial (HBE) cells with ΔF508 CFTR mutation, affects nearly a dozen porosome proteins including CFTR. Therefore, we hypothesized that the introduction of normal functional porosomes carrying wild type CFTR into the cell plasma membrane of CF cells would rescue from all forms of CF. Air liquid interface (ALI) 3D differentiated HBE WT-CFTR cells and ΔF508-CFTR CF HBE cell cultures mimicking normal lung physiology, responding to CFTR inhibitors and CF corrector and modulator drugs Tezacaftor, Ivacaftor and TRIKAFTA, was used in the study. Introduction of functional porosome complexes obtained from WT-CFTR HBE cells into the plasma membrane (PM) of ΔF508-CFTR CF cells, was demonstrated by an increase in PM-associated CFTR using Magnify expansion microscopy. Mucin secretion assays demonstrate porosome reconstitution to restore mucin secretion more than twice as effectively as TRIKAFTA. These results are further supported by preliminary nasal potential different studies in ΔF508 mice, where treatment of the nasal passage with porosome isolates from WT-CFTR HBE cells, restore chloride secretion in the nasal passage of mice, a further validation of the highly effective porosome reconstitution therapy for CF.

## INTRODUCTION

Human airway epithelium is coated with a thin film of mucus, composed primarily of mucin MUC5AC and MUC5B [1], that is propelled and cleared via ciliary motion. This ciliary clearance of mucus, assists in keeping the airway moist, lubricated and clean, free from infection. The major issue in cystic fibrosis (CF), is increased mucin viscosity and the inability of cilia to propel and clear the mucin, leading to its stagnation and infection. Studies report that in CF patients, there is loss of both MUC5AC and MUC5B mucin in sputum [2]. CF patients exhibit a nearly 70% decrease in MUC5B and up to a 93% decrease in MUC5AC in their sputum compared to subjects without any lung disease [2]. Our studies in Calu-3 cells, a human airway epithelial cell line, also demonstrate loss of both chloride and mucus secretion [3] as demonstrated following exposure to either CFTR inhibitors 172 or GlyH101. Porosomes are secretory portals at the cell plasma membrane where secretory vesicles transiently dock and fuse to expel a precise amount of intra-vesicular contents from the cell during secretion [4,5]. Since proteome of the porosome secretory machinery in the human airways epithelia is composed of the CFTR protein [6], implicates CFTR in porosome-mediated mucus secretion.

In a recent study [7], ALI differentiated WT-CFTR Human Bronchial Epithelial (HBE) 3D cell cultures and ΔF508-CFTR CF HBE 3D cultures that respond to CFTR inhibitors [3,6], also respond to the CF corrector/modulator drugs Tezacaftor and Ivacaftor [7]. This ALI 3D model was found to closely mimick the physiological state of the human airway epithelium, and hence has greatly helped in assessing CF treatments and therapy prior to human clinical trials. Western blot analysis of porosomes isolated from WT-CFTR HBE expanded cell cultures and ΔF508-CFTR CF HBE expanded cell cultures, demonstrated a loss of the t-SNARE protein SNAP-23 in the ΔF508-CFTR cells [Figure 1a, Ref. 7]. Similarly, mass spectrometry of porosomes isolated from WT-CFTR HBE Cells and ΔF508-CFTR CF HBE cells, demonstrate a varying loss or gain of several porosome proteins [7]. These studies demonstrate that mutation in CFTR impacts other proteins within the porosome secretory complex besides CFTR. Therefore, expression of just the normal CFTR protein in treating CF is inadequate in treating the disease. In the porosome complex composed of around 34 proteins, it would be nearly impossible to replace the mutated CFTR protein with the normal functional wild type protein. Earlier attempts therefore using gene therapy, have not been very effective in treating CF. Hence, the currently available successful CF treatments rely on the use of small molecular modulator products to correct the mutated CFTR function by varying degrees (up to 13-14%) in CF patients. None of the current CF therapies however, address the disease as a defect in the mucus secretory machinery of the cell. Therefore, to address this issue of multi-protein malfunction in CF, in addition to correcting the CFTR protein, and to be able to treat all forms of CFTR mutations, the porosome reconstitution therapy [Figure 1b, Ref.7] was adopted. In the earlier study [7], porosome reconstitution in ΔF508-CFTR HBE CF cells was able to greatly increase MUC5B and MUC5AC secretions. The two CF drugs Tezacaftor and Lvacaftor were also used in the study to compare effectiveness in restoring normal MUC5B and MUC5AC secretions. Results from the study [7] show that on average, porosome reconstitution was able to restore by >40% the secretion of MUC5B in ΔF508-CFTR cells. Similarly, in ΔF508-CFTR CF cells, a 9% and 11% increase in MUC5AC secretion is observed in the presence of Lvacaftor and Tezacaftor respectively, only on day 2 following exposure to the drugs, while porosome-reconstitution in the same period demonstrated around a 44% increase, a near four-fold greater than the two CF drugs Lvacaftor and Tezacaftor.

**Figure 1.**
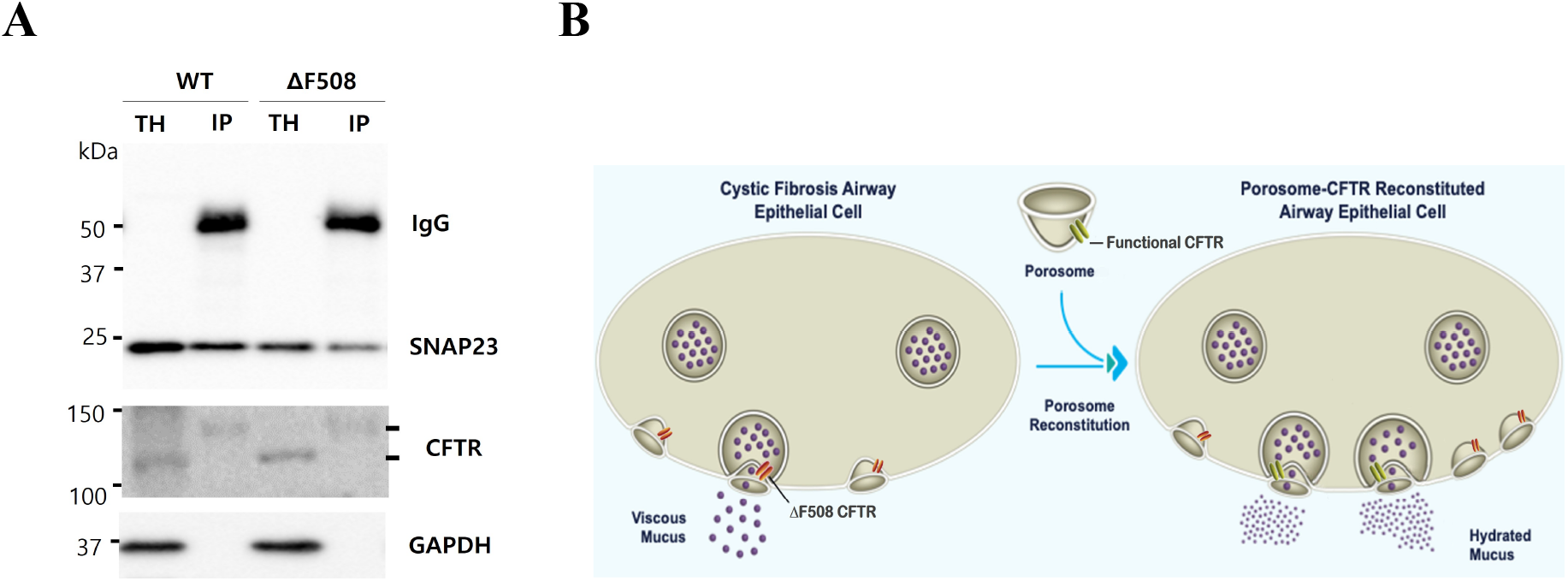
In human bronchial epithelial (HBE) cells with ΔF508-CFTR mutation, there is a loss of SNAP-23 protein in the porosome complex, which could be replaced by the introduction or reconstitution of porosome complexes obtained from WT-CFTR HBE cells into the plasma membrane of the ΔF508-CFTR CF cells. **(A)** Porosome isolated from WT-CFTR HBE cells and ΔF508-CFTR CF HBE cells using SNAP 23 antibody demonstrate a decrease in the presence of SNAP-23 protein both in total homogenate (TH) and in immunoisolated (IP) porosomes from ΔF508-CFTR CF HBE cells [7]. These results suggest that altered CFTR protein within the porosome complex as a consequence of mutation in the CFTR gene alters other proteins within the porosome complex, altering porosome function. However, the total amount of the CFTR protein in the porosome of ΔF508-CFTR CF HBE cells do not appear to change and is present in both the WT and ΔF508 cells in the high molecular glycosylated state. In TH of WT and ΔF508 cells, both the non-glycosylated low molecular weight CFTR and the glycosylated higher molecular weight CFTR forms are present. **(B)** Schematic representation of porosome-reconstitution therapy for cystic fibrosis (CF). Functional normal CFTR porosomes obtained from WT-CFTR HBE cells, will rescue from CF by normalizing mucus secretion [7].

To further test the potency and efficacy of the porosome reconstitution therapy in treating CF, and to understand the expression and distribution of the porosomes following their reconstitution, the current study was performed. In this study, porosome reconstitution therapy was also compared with the current most effective drug TRIKAFTA [8]. ALI differentiated 3D cultures of ΔF508-CFTR HBE CF cells were reconstituted with different amounts of isolated porosomes, and both the expression and distribution of porosome proteins following reconstitution was assessed using Magnify Expansion Microscopy (Magnify ExM) [9, 10]. To determine secretory function, MUC5AC secretion was assessed in the 3D cultures of ΔF508-CFTR HBE CF cells following porosome reconstitution or in the presence of the current CF drug TRIKAFTA. Results from the current study show great promise of the porosome-reconstitution therapy in treating CF including all forms of CFTR mutations.

## 2. MATERIALS AND METHODS

### 2.1 Human Bronchial Epithelial Cell Line WT-CFTR and ΔF508-CFTR ***Cultures***

WT-CFTR Human Bronchial Epithelial (HBE) cell line (CFBE41o-6.2) and experimental ΔF508-CFTR HBE CF cell line (CFBE41o) a ΔF508del (-/-) homozygous, was obtained from Sigma (Temecula, CA 92590, USA). Cells were stored, handled and cultured using established published procedures and protocols outlined in the data sheet. Cells were cultured using Fibronectin/Collagen/BSA ECM mixture-coated petri dishes (10µg/mL Human Fibronectin, Sigma Cat. No. F2006; 100µg/mL BSA, Sigma Cat.No.126575; 30µg/mL PureCol, Sigma Cat No. 5006; α-MEM Medium, Sigma Cat. No. M2279) with α-MEM Medium supplemented with 2mM L-Glutamine (Gibco Cat. No. 25030-081, Grand Island, NY 14072, USA); 10% fetal bovine serum (Sigma Cat. No. ES-009-B); and 100 U/ml Penicillin and 100 µg/ml Streptomycin (Gibco Cat. No. 15140-122). Cultures were incubated at 37°C in 95% air/5% CO_2_ atmosphere.

### 2.2 Differentiated Human Bronchial Epithelial 3D Cell Cultures

Air liquid interface (ALI) differentiated WT-CFTR and ΔF508-CFTR CF HBE cell cultures mimicking normal lung physiology, were established using a minor modification of our published procedure [7]. These ALI 3D cultures respond to CFTR inhibitors [3] and the CF corrector/modulator drugs Tezacaftor and Ivacaftor [7]. To establish ALI 3D cultures, cells were seeded in sterile ECM mixture-coated transwell inserts of 12 mm diameter with 0.4 µm pore size (Corning Cat. No. Costar 3460, Kennebunk. ME 04043 USA) with a seeding density of 2 × 10^5^ cells/insert. Media of apical and basolateral region of transwell were changed every other day, 500µL on the apical side and 1mL on the basolateral side. At confluency on day 7, cells were raised to ALI condition without apical medium. On day 21 after switching to ALI, cultures were treated with either TRIKAFTA composed of 20 µM Tezacaftor, 3 µM Elexacaftor, and 5 µM Ivacaftor (Selleck Chemicals, Houston, TX 77014 USA) or immunoisolated porosomes (0.2 µg/mL, 0.5 µg/mL, 1µg/mL).

### 2.3 Porosome Isolation and Reconstitution into ΔF508-CFTR HBE CF Cells & ΔF508 Mice Nasal Epithelia

WT-CFTR HBE cell line (CFBE41o-6.2) and experimental ΔF508-CFTR HBE CF cell line (CFBE41o) a ΔF508del (-/-) homozygous were used to isolate porosomes for proteome analysis and reconstitution. SNAP-23 specific antibody (Abcam Cat. No. AB3340, Cambridge, UK) was used to immunoisolate porosomes from solubilized cells. Protein in all fractions was estimated using BCA Protein assay Kit (ThermoFisher Cat. No. 23227, Rockford, IL 61101, USA). SNAP-23 specific antibody-crosslinked to protein A/G Magnetic-agarose (ThermoFisher Cat. No. 78609) was used. To reduce the antibody contamination in eluted protein solution, the antibody was chemically crosslinked to the agarose-magnetic beads. Briefly, the beads were resuspended in dilution buffer (1:1 ratio, 1mg/mL BSA in PBS) for 10 min at 4°C, centrifuged for 1 min at 14,000 rpm in a bench top centrifuge, and the supernatant aspirated. SNAP-23 antibody (1µg/mL) in dilution buffer was added to the beads at 1:1 ratio and mixed gently for 1hr at 4°C. The beads were then washed twice with 10 volumes of dilution buffer. Dimethyl pimelimidate solution (DMP, 13mg/ml, Sigma Cat. No. D8388) in wash buffer (0.2 M triethanolamine in PBS, Sigma Cat. No. 90279) was added to the SNAP23 antibody conjugated beads (1:1 ratio) and resuspended for 30 min at room temperature (RT). The beads were then washed with wash buffer three time. (30 min/wash at RT), and resuspended in quenching buffer (50mM ethanolamine in PBS, Sigma Cat. No. E0135) for 5 min at RT and washed with PBS twice. To remove excess unlinked antibody, the beads were washed with 1 M glycine pH 3, twice (10min/wash at RT). Prior to use for immunoprecipitation, the beads were washed in PBS-TWEEN buffer three times. Cells were solubilized in Triton/Lubrol solubilization buffer (0.5% Lubrol; 1 mM benzamidine; 5 mM ATP; 5 mM EDTA; 0.5%Triton X-100, in PBS), supplemented with protease inhibitor mix (Sigma, St. Louis, MO). SNAP-23 antibody-crosslinked to the protein A/G Magnetic-agarose was incubated with the solubilized cell lysates for 16h at 4°C followed by washing with wash buffer (500 mM NaCl, 10 mM TRIS, 2 mM EDTA, pH 7.5). The immune-isolated porosomes associated with the immuno-agarose beads were dissociated and eluted using pH 3.0 PBS solution, and the eluted sample was immediately returned to neutral pH prior to Western Blot analysis and reconstitution assays. Porosome reconstitution into ΔF508-CFTR CF HBE cells was achieved by exposing 0.2 µg/mL, 0.5 µg/mL or 1µg/mL porosomes isolated from WT-CFTR HBE cells. Similarly, 4µg/mL porosomes isolated from WT-CFTR HBE cells were used to reconstitute into the nasal epithelia of mice. Porosome reconstitution experiments were conducted using a B6.129 Cftrtm1KthTg(FABPCFTR)1Jaw/Cwr mice. They are a ΔF508 line backcrossed onto a C57bl/6 background strain. The mice strain express CFTR in the gut driven by the FABP promoter to avoid intestinal blockage. Porosomes obtained from WT-CFTR HBE cells were used to treat the nasal passage of the CF mice and the vehicle (PBS pH7.5) alone was used as control. Experimental animals were treated using four micrograms of the isolated porosome complexes in PBS pH 7.5 using a nasal catheter and nasal potential difference (PD) was measured after 24h post porosome reconstitution, to determine CFTR ion channel activity. In this experiment, nasal potential difference is used to measure the voltage across the nasal epithelium, which results from trans-epithelial ion transport reflecting in part, CFTR function.

### 2.4 ELISA

At different time point (1day, 2day) following treatment with TRIKAFTA or isolated porosomes, the apical surface, basal surface, or both, of each ALI culture, was gently washed and aspirated using 500uL fresh PBS. These washes were stored at 4°C for ELISA assays for MUC5AC. To quantify the mucin secretion, 100uL of each wash was added to the ELISA plate. MUC5AC ELISA Kit (MyBioSource Cat. No. MBS701926, San Diego, CA 92195, USA) were used according to the manufacturer’s instructions. Optical density was determined at 450nm using a BioTek Synergy HT microplate reader (BioTek, Winooski, VT 05404, USA), and the mucins quantified.

### 2.5 Magnify Expansion Microscopy

#### Pre-expansion Immunostaining and Imaging

Human Bronchial Epithelial (HBE) cells, reconstituted with porosomes, were grown in air-liquid interface (ALI) cultures to model human lung tissue for cystic fibrosis (CF) studies. Fixed wild-type and CFTR ΔF508-mutated cells, treated with porosome concentrations (0, 0.2, 0.5, and 1 µg), were mounted on glass slides. Cells were incubated with a staining buffer (2X SSC, 0.2% Tween-20, 1X PBS) containing primary antibodies (1:500 dilution): mouse anti-CFTR (Invitrogen; MA1-935) and rabbit anti-SNAP23 (Invitrogen; PA5-28936) for 45 minutes to 1 hour at room temperature.

Following two 15-minute washes with the same buffer, secondary antibodies (1:500 dilution) were applied: donkey anti-mouse IgG (H+L) DyLight 488 (Invitrogen; SA5-10166) and goat anti-rabbit IgG (H+L) DyLight 500 (Invitrogen; SA5-10033), along with DAPI (1:1000). The samples were washed again for 15 minutes and imaged using a Keyence BZ-X810 microscope.

#### In Situ Polymerization of CF Cells using Magnify Protocol

Monomer solution was prepared (4% DMAA (v/v), 34% sodium acrylate (w/v), 10% acrylamide (w/v), 0.01% bis-acrylamide (w/v), 1% NaCl (w/v) in 1X PBS). Before gelation, 0.25% APS, 0.1% TEMED, and 0.1% methacrolein were added. The solution was vortexed, and cells (membranes facing upward) were incubated with the gelling solution at 4°C for 15 minutes to ensure diffusion. A gelling chamber was created using a no. 1.5 coverslip, and the samples were incubated overnight at 37°C in a humidified container to complete gelation. Following polymerization, the upper coverslip was removed, and excess gel was trimmed. The samples were homogenized in 10% SDS, 8M urea, 25 mM EDTA, and 2X PBS (pH 8.4) for 45 minutes at 95°C. The plastic membranes were carefully peeled away, and the gels were washed three times with 0.1% C12E10/1X PBS to remove residual SDS.

#### Immunostaining of Gel-embedded Samples

Post-homogenization, the samples were stained by incubating with primary antibodies (1:250 dilution) in a staining buffer (2X SSC, 1X PBS): mouse anti-CFTR (Invitrogen; MA1-935) and rabbit anti-SNAP23 (Invitrogen; PA5-28936) for 1.5 hours at room temperature or 1 hour at 37°C. After three washes, secondary antibodies (1:500) donkey anti-mouse IgG (H+L) DyLight 488 and goat anti-rabbit IgG (H+L) DyLight 500 were applied along with DAPI (1:1000). The samples were washed again three times before imaging.

#### Imaging Modality

Wide-field fluorescence imaging was performed using a Keyence BZ-X810 microscope equipped with the following objectives: BZ-PF04P Plan Fluorite 4× (0.2 NA), BZ-PF10P Plan Fluorite 10× (0.4 NA), and BZ-PF20LP Plan Fluorite 20× (0.5 NA). The microscope system allowed for high-quality imaging of stained samples at various magnifications, providing detailed visualization of cellular structures. Images were captured and processed using the Keyence analysis software, ensuring accurate and reliable results.

Magnify imaging was performed using a Nikon Eclipse Ti2 epifluorescence microscope equipped with a CSU-W1 spinning disk confocal module and an Andor 4.2 Zyla sCMOS camera. The system was controlled by NIS-Elements AR 5.21.03 64-bit software. Imaging was carried out using the following Nikon objectives: CFI Plan Apo Lambda 4× (0.2 NA), CFI Plan Apo Lambda 10× (0.45 NA), CFI Apo LWD Lambda S 20×WI (0.95 NA), CFI Apo LWD Lambda S 40×WI (1.15 NA), and CFI Plan Apo Lambda 60×Oil (1.4 NA).

## 3. RESULTS AND DISCUSSION

Magnify expansion immunocytochemistry [Figure 2] demonstrate that in ΔF508-CFTR CF HBE cells, the CFTR protein fails to traffic to the cell plasma membrane as previously reported [11, 12], and is distributed in the cytosol. Increasing amounts of porosome reconstitution, demonstrates association of both the porosome proteins SNAP-23 and CFTR at the cell plasma membrane of ΔF508-CFTR CF HBE cells [Figure 2]. Transmission electron microscopy demonstrates that the overall morphology of ΔF508-CFTR CF HBE cells following porosome reconstitution, return to those present in WT CFTR HBE cells (data not shown). Parallel experiments carried out in 3D differentiated ΔF508-CFTR CF HBE cultures, demonstrate a more than 2-fold increase (P<0.05) in MUC5AC secretion in porosome reconstituted cells, compared to TRIKAFTA treatment in the same period. These results in cells were further supported by preliminary nasal potential different studies in B6.129 *Cftr*^*tm1Kth*^Tg(FABPCFTR)1Jaw/Cwr ΔF508 mice. In these mice studies, treatment of the nasal passage with porosome obtained from WT-CFTR HBE cells rescues chloride secretion in the nasal passage, even after 4 days following treatment, demonstrating a less frequent requirement of the porosome treatment as opposed to the daily requirement of TRIKAFTA. Additionally, these animal studies reflect cross-species efficacy and immune tolerance in the porosome reconstitution, demonstrating great promise in the porosome reconstitution therapy for the treatment of all forms of CF.

**Figure 2.**
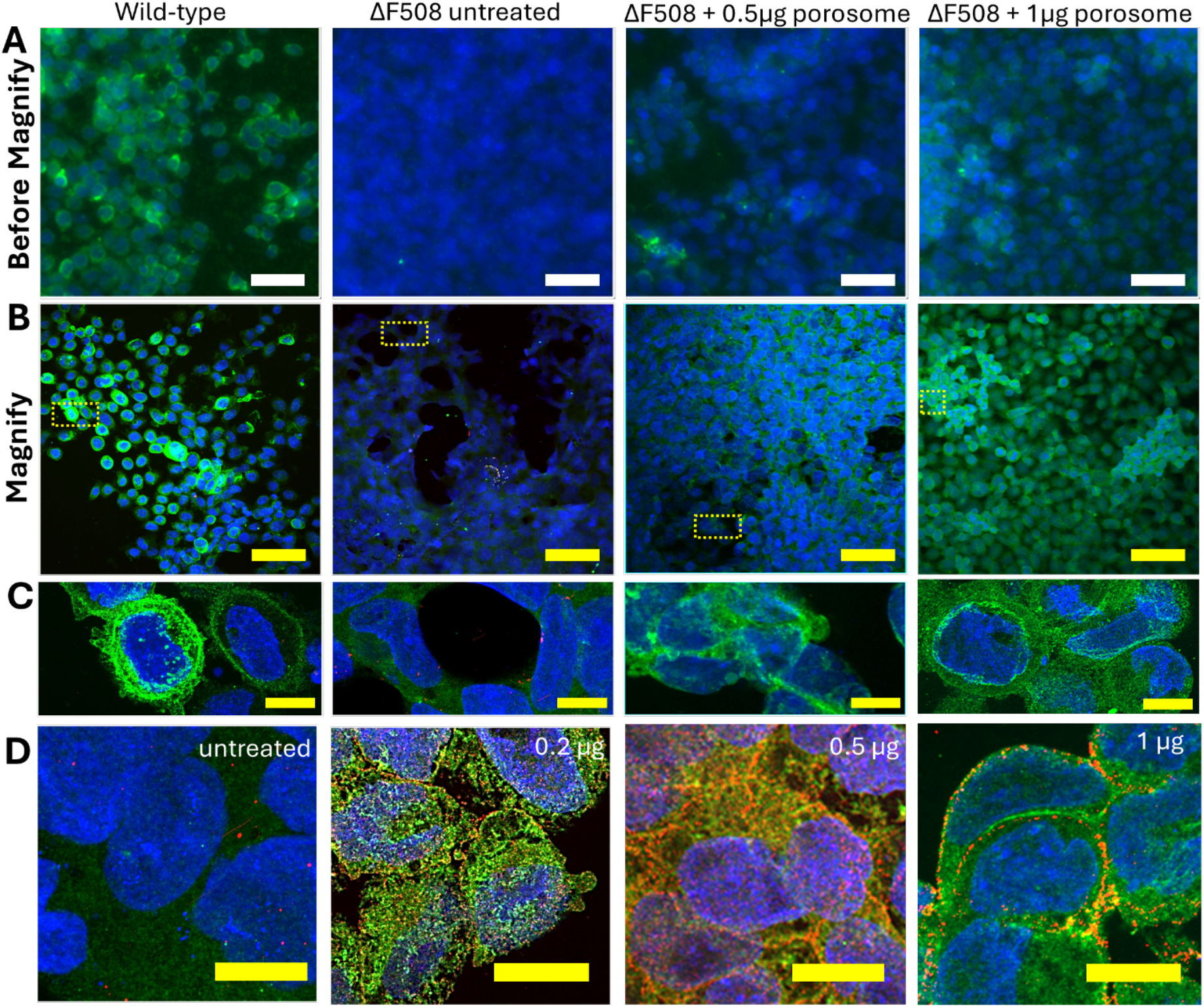
Wide-field and Magnify-processed images of Human Bronchial Epithelial (HBE) cells showing porosome treatment restores expression and localizations of CFTR and SNAP23 in CFTR ΔF508-mutated cells. **(A)** Wide-field fluorescence images of untreated HBE cells (left) and HBE cells with CFTR ΔF508 mutation treated with increasing concentrations of porosomes (0.5, and 1 µg). The scale bar represents 50 µm. DAPI (blue) stains the nuclei, and CFTR (green) highlights the CFTR protein. (**B)** Magnify-processed images of the same conditions as in Panel A, showing enhanced resolution and detail of CFTR expression. The scale bar represents 50 µm. (**C)** Zoomed-in images of the areas indicated by yellow-dotted boxes in Panel B. These images provide a closer view of CFTR expression in the HBE cells. The scale bar represents 5 µm. **(D)** Magnify-processed images of CFTR ΔF508-mutated HBE cells treated with increasing doses of porosomes (0.2, 0.5, and 1 µg). This panel shows DAPI (blue), CFTR (green), and SNAP23 (red), illustrating the expression and localization of both proteins in response to porosome treatment. The scale bar represents 5 µm in biological scale. **B, C and D:** The expansion factor is 3.8× (1× PBS).

**Figure 3.**
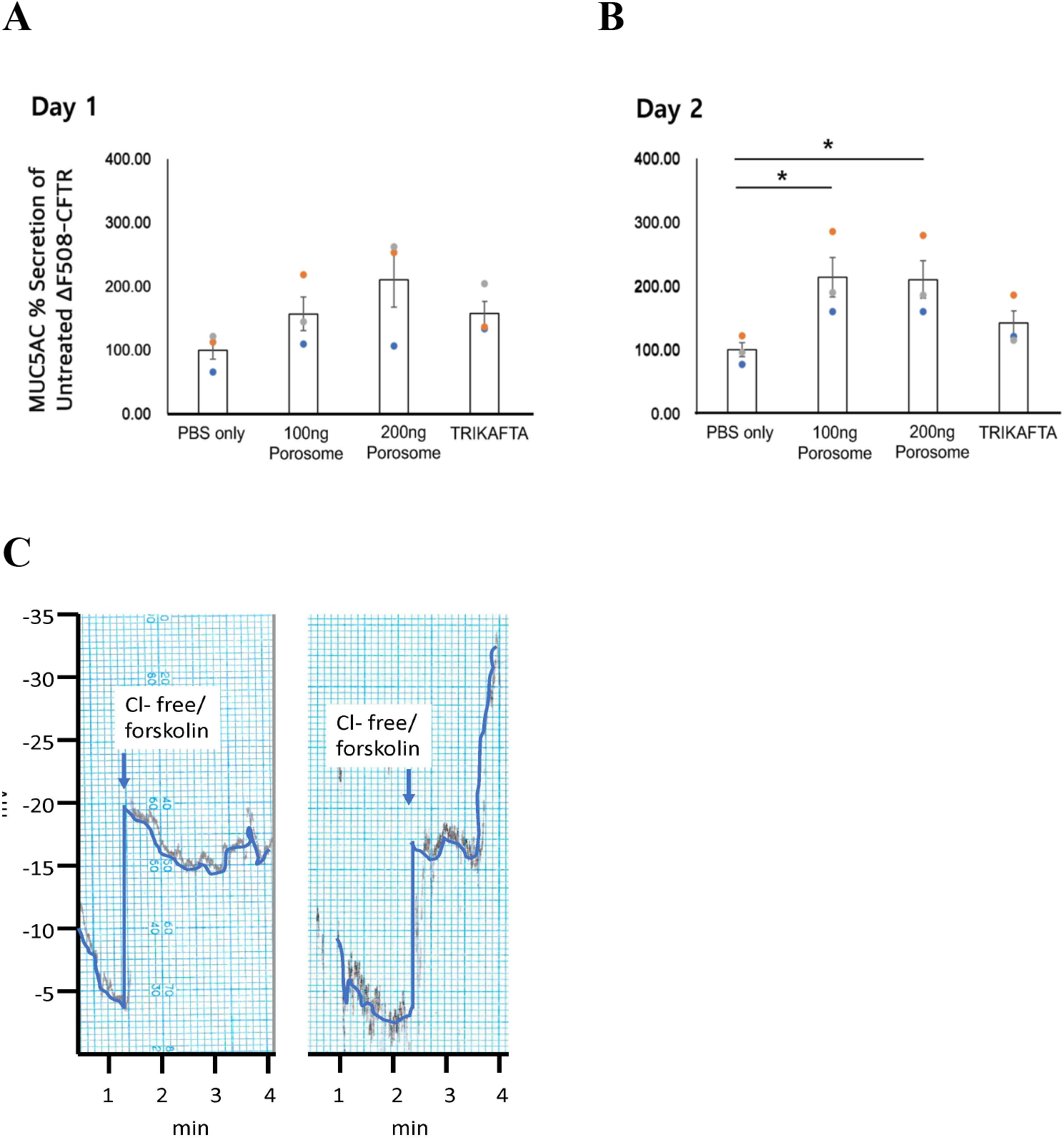
Functional reconstitution of normal CFTR-associated porosomes obtained from WT-CFTR HBE cells into differentiated 3D ΔF508-CFTR CF HBE cells in culture, as well as reconstitution in the nasal passage of ΔF508-CFTR CF mice, potentiates mucus and chloride secretion, resulting in rescue from cystic fibrosis. **(A, B)** In ΔF508-CFTR CF cells, a significant (P<0.05) and much greater increase (>2-fold over TRIKAFTA) in MUC5AC secretion is observed following porosome-reconstitution, compared to TRIKAFTA in the same period. **(C)** Preliminary, nasal potential difference assay following porosome reconstitution in nasal passage of CF mice. Porosome obtained from WT-CFTR HBE cells rescues chloride secretion in nasal passage of ΔF508-CFTR CF mice restores CFTR function. Nasal potential difference assay of mice treated with carrier **(Left)** or 4 µg porosome **(Right)** for two consecutive days. Mice were analyzed 24 hours after the last treatment. Mice are first treated with amiloride containing Ringer’s to block sodium transport, and then are switched to chloride-free Ringer’s with forskolin (10 µM) and amiloride (10 µM) to drive chloride secretion. Upward inflection indicates CFTR function. Mice respond to forskolin stimulation even after 4 days following porosome reconstitution, demonstrating functional stability of the reconstitution.

As previously reported [7], ALI certified normal human bronchial epithelial cells using an established rotating bioreactor for scalable culture and differentiation of the human respiratory epithelium [57] could be utilized to obtain the biologic for commercial use in therapy. Since the human airway epithelium is the primary target for all inhaled air-borne substances, and porosomes being nano size (∼100 nm) particles that are retained at the cell plasma membrane, their delivery to the lung epithelia would be the ideal therapeutic approach for CF.

## Contributions

The idea for the use of porosome-reconstitution therapy was developed at Porosome Therapeutics, Inc. The research design was developed by B.P.J. and conducted by W-J.C. Magnify and electron microscopy were performed in the laboratory of YZ and DJT respectively. PD studies in mice were performed in the CF mice core at Case Western Reserve University as paid contract from Porosome Therapeutics, Inc. B.P.J. wrote the paper. All authors participated in critical reading and discussion of the manuscript.

## Acknowledgements

Work presented in this article was supported by the Viron Molecular Medicine Institute and Porosome Therapeutics, Inc., Boston, MA.

